# Hippocampal β-adrenergic system modulates recognition memory reconsolidation

**DOI:** 10.1101/2022.05.16.492176

**Authors:** Gustavo Balbinot, Josué Haubrich

## Abstract

Targeting reconsolidation with propranolol, a blocker of β-adrenergic receptors (β-ARs), emerged as a potential treatment for maladaptive memories such as those involved in posttraumatic stress disorder (PTSD). Reconsolidation targeting treatments for PTSD are becoming a common practice in the clinic and it is important to unveil any side effects upon ‘non-targeted’ memories. While previous studies have focused on propranolol’s effects on the reconsolidation of emotional/distressful memories, the present study asked whether propranolol is involved in the reconsolidation of recognition memories - by assessing its effects on distinct memory components and the role of the hippocampus. Rats performed an object recognition (OR) task where they were exposed to different objects: A and B presented during the sample phase; A and C presented during the reactivation phase; and D in combination of either A, B, or C during a final test. Intra-hippocampal injections of propranolol (5 µg or 10 µg) were conducted immediately after the reactivation session. Propranolol infusions consistently impaired the addition of novel information to the previously consolidated memory trace regardless of dose, and the retention of familiar objects was not affected. Higher doses of propranolol also hindered memory of a familiar object that was not presented during the reactivation session, but was previously placed at the same location where novel information was presented during reactivation. The present results shed light on the role of β-ARs on the reconsolidation of different memory components and argue for the need for further studies examining possible recognition memory deficits following propranolol treatment.

**Graphical abstract:** 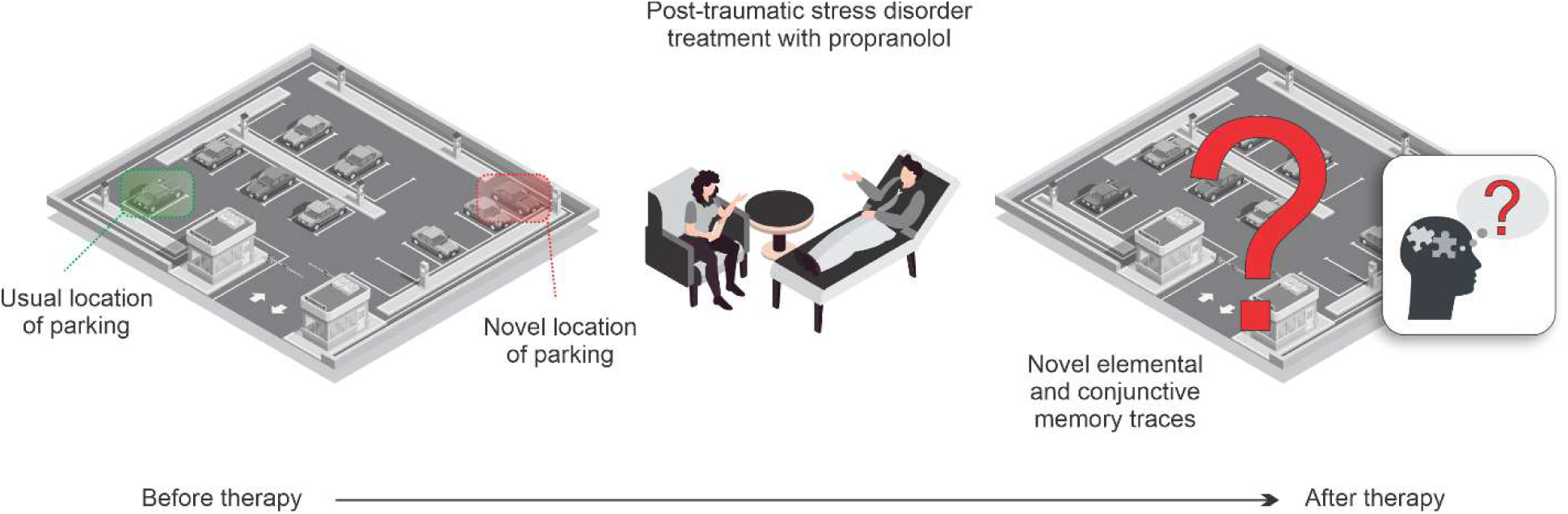

Post-traumatic stress disorder is a chronic mental health condition, which may develop following direct or indirect exposure to a traumatic event. The administration of propranolol to individuals affected by this disorder before the reactivation of the trauma-related memory may diminish the symptoms of this mental condition. Here, we show that such treatment may have effects on non-targeted memories, other than the fear/distressful memory. In a series of experiments in rodents, we show that intra-hippocampal infusion of propranolol immediately after recalling and updating a recognition memory trace hampers the reconsolidation of the initial recognition memory trace during recall. This may lead to difficulties in recalling recent events related to declarative memories. In a high dose, propranolol treatment may also affect the conjunctive component (association between multiple elements) of the memory trace – in addition to the effect on the elemental component. This may lead, for example, to difficulties in locating a parked car in a non-usual location after the post-stress traumatic stress disorder therapy with propranolol.

**Highlights:**

- We established the role of the hippocampal β-adrenergic system in the reconsolidation of recognition memories.
- Propranolol treatment may impair the updating of recognition memory traces.
- High doses of propranolol may disrupt both elemental and conjunctive components of memory.
- Clinical treatment with high doses of propranolol for post-traumatic stress disorder may unintentionally affect non-pathological components of memories.

## 1. Introduction

Reactivation of a previously consolidated memory can bring it to a labile state, allowing the updating of its content with novel information, a process termed reconsolidation (Nader Karim et al., 2000). During reconsolidation, memories become sensitive to modifications, providing the opportunity to alter unwanted memories, such as in posttraumatic stress disorder (PTSD) (Haubrich et al., 2020; Schwabe et al., 2012). The β-adrenergic receptor (β-AR) blocker propranolol emerged as a potential pharmacological tool for targeting reconsolidation in PTSD, as such, experimental and clinical studies have evaluated its effects on the reconsolidation of fear or distressful memories (Giustino et al., 2016; Roullet et al., 2021; Schwabe et al., 2012; Weymar et al., 2010). Given that reconsolidation-based treatments for maladaptive memories are potentially becoming psychologists’ common best practice, any side effects upon non-targeted memories must be unveiled.

Reconsolidation-blockade treatments aim to attenuate maladaptive memories originating from prior experiences (Phelps and Hofmann, 2019). Often, these experiences involve the co-occurrence of adverse and neutral outcomes in a defined location in time and space (*i.e*., in a context). Thus, memories of these events encompass a collection of independent elemental features bound together into a conjunctive representation that encodes the co-occurrence of these elements, as well as their specific location (Rudy et al., 2004). It is unclear which components of such memories – elemental or conjunctive – are most affected by reconsolidation-targeted treatments. The object recognition (OR) task is useful in this regard because both elemental and conjunctive components of memory can be assessed (Cohen et al., 2013). The OR memory encodes information about features of different objects and of the context where the objects are in (elemental representations). It also associates specific objects with specific locations in the context (conjunctive representation). Moreover, recognition memories are of fundamental importance for distinguishing familiar and novel stimuli and can be viewed as a model for episodic-like memory in rodents (Antunes and Biala, 2012; Dere et al., 2005).

Reconsolidation of recognition memories is hippocampal protein synthesis-dependent, and the hippocampus is likely to mediate the addition of new information to a previously consolidated memory trace (Rossato et al., 2007). It has been previously reported that hippocampal β-ARs modulate BDNF expression and the consolidation of OR memories (Furini et al., 2010), and that systemic post-reactivation propranolol treatment disrupts reconsolidation of OR memory (Villain et al., 2016). To date, however, there is no study on the possible effects of intra-hippocampal infusions of propranolol on the reconsolidation of recognition memories. It is also not known which component of OR memory is affected by this intra-hippocampal propranolol infusion. The present study aimed to evaluate the effects of hippocampal β-ARs blockade by propranolol on the reconsolidation of recognition memories and to determine which memory component is affected. We hypothesized that intra-hippocampal propranolol infusions would affect the reconsolidation of a novel recognition memory trace by selectively disrupting the incorporation of novel information.

## 2. Materials and methods

### Ethical statement

All the animal manipulation and procedures described in this manuscript followed the local regulatory laws and the 3R’s principles to minimize the number of animals involved in the research. All experimental procedures were approved by the Institutional Animal Care and Use Committee of the institution (CEUA/UFRN; protocol number 050/2015).

### 2.1. Animals

Male Wistar rats (*Rattus novergicus*, 3 months old, 300-350 g) were housed in isolated ventilation cages (5 animals/cage; standard polysulphone box; 49 cm x 34 cm x 16 cm; Alesco®, SP/BR), in a 12h/12h light/dark cycle with controlled temperature (21ºC ± 2ºC), and received food and water *ad libitum*. A total of 130 Wistar rats were used in two separate experiments (obtained from CREAL/UFPB/BR). Two animals were excluded due to sickness and two animals were excluded from analysis due to minimum exploration time criteria (15 s) (Cohen and Stackman Jr., 2015).

### 2.2. Experimental protocols

#### 2.2.1. Object recognition (OR) task

Before the novel OR task, the animals were habituated to the arena for 4 days (20 minutes of habituation per day). The arena is a square box (60 cm x 60 cm) with lighting conditions kept at ≈ 40 Lumix and the room temperature controlled (≈ 24°C). In the sample phase, animals were exposed to two objects (A and B) for 10 minutes (Bevins and Besheer, 2006). After a 24 hours interval, the animals were re-exposed to one previously presented object (A) and to a novel object (C) for 5 minutes (reactivation phase). Immediately after the reactivation phase, we performed bilateral intra-hippocampal injections of 1µL of saline or propranolol (5 µg/µL or 10 µg/µL). The test phase took place 24 h later and animals were assigned into one of three groups where a novel object (D) was presented along with one of the familiar objects (A, B, or C), as follows: objects A and D, objects D and B, or objects D and C (Figure 1a). Exploration was defined as sniffing or touching the object with the nose and/or forepaws. Touching the objects with forepaws while the nose was not directed to the objects, turning around, or sitting on the object were not considered object exploration. A recognition index calculated for each animal was expressed as (T_X_ or T_Y_)/(T_X_ + T_Y_) (T_X_ = time spent exploring the familiar object; T_Y_ = time spent exploring the novel object) (Cohen and Stackman Jr., 2015). The amount of time spent exploring each object was recorded by a video camera (CPF55; Pyxel Eletronics®, PR/BR). A total exploration minimum criterium was defined *a priori* (15 s) (Cohen and Stackman Jr., 2015). A trained observer blinded to the treatment manually scored exploration times. Intact recognition memory in the test phase is reflected in a discrimination score higher than 0.5, which implies greater exploration of the novel object.

**Figure 1.**
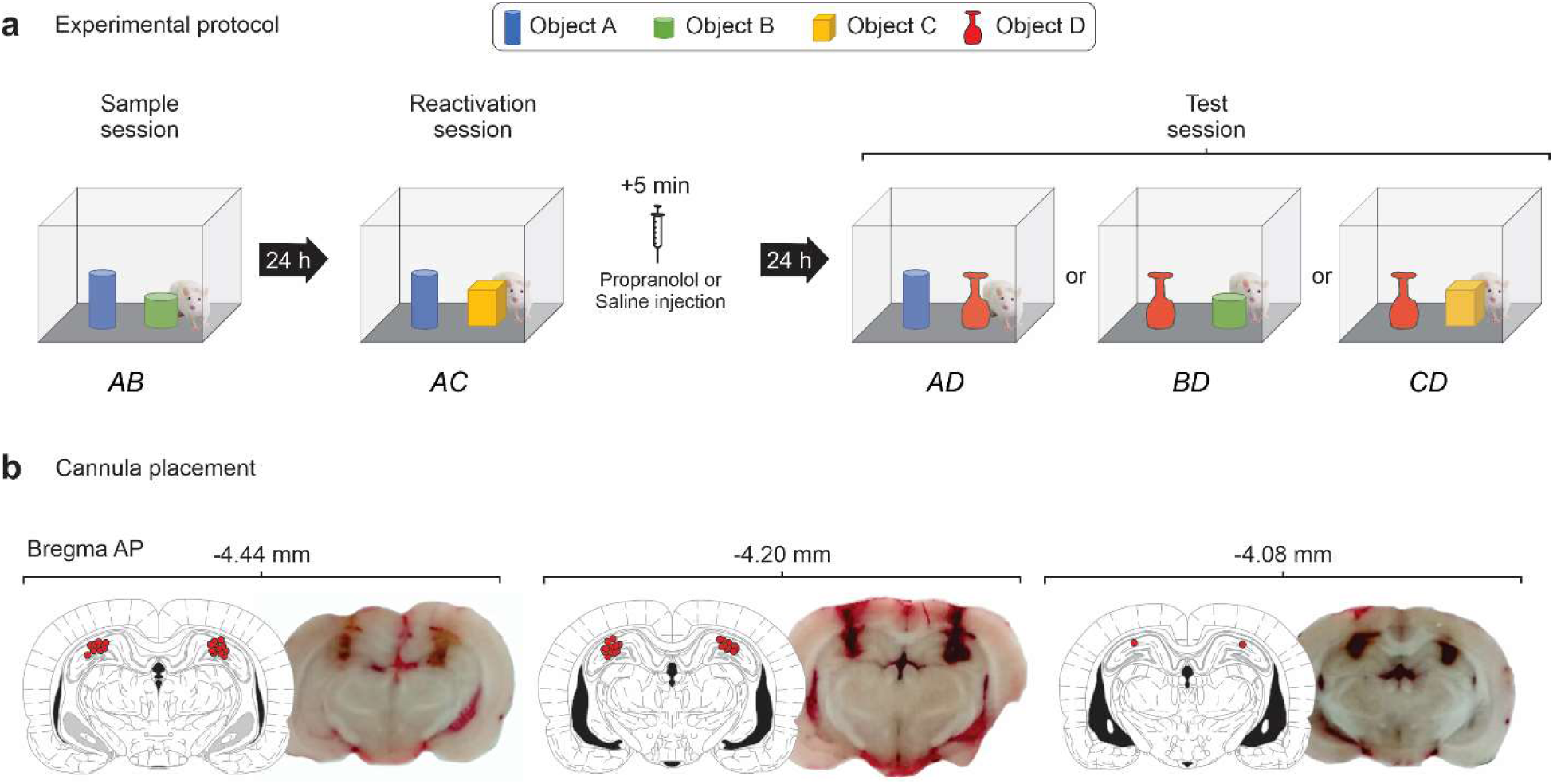
Experimental protocol and cannula placement. (**a**) Rats with infusion cannula implanted in the CA1 region of the dorsal hippocampus were exposed to two objects (A and B) for 10 min (Sample phase; Day 1). Twenty-four hours later the animals were exposed to a familiar object A in combination with a novel object (C) (Reactivation phase; Day 2). Rats were randomly assigned to one out of three different groups and immediately after that received bilateral intra-CA1 infusions of saline (1 μL/side) or propranolol (5 μg/side or 10 μg/side). Twenty-four hours later animals were submitted to a 5-min-long test phase (Test phase; Day 3) in the presence of different combinations of objects, as follows: (Group 1) Object A + Object D; (Group 2) Object B + Object D; (Group 3) Object C + Object D, where D was a novel object. (**b**) Cannula placement in the dorsal hippocampus. Schematic of cannula placements for a representative sample of rats (n = 27) displayed on a plate from Paxinos & Watson (1998). Each red circle represents the core of each infusion. Cannula placement was verified by the injection of methylene blue (1 μL/side) before sacrifice. Photomicrograph showing three methylene blue-stained coronal sections from three different brains of rats with representative cannula placements in the dorsal hippocampus. Cannula positioning varied from AP -4.44 to AP -4.06 relative to bregma. Note that the drug infusion reached the dorsal hippocampus mainly at CA1 region, but extended to further hippocampal areas.

#### 2.2.2. surgical cannula placement

One week before behavioral testing, rats were implanted with stainless steel guide cannulas aimed at the dorsal hippocampus (Insight®, SP/BR; AP -4.2, ML ±3.0, DV -1.2) (Paxinos and Watson, 2007). All surgical procedures were conducted with the animals under ketamine and xylazine anesthesia (intra-peritoneal injections; 75 mg/Kg and 10 mg/Kg, respectively). After surgery, the animals received analgesics for two days (sodium metamizole; 0.1 mL/animal every 4 hours; Neo Química®, Brazil). To allow a good recovery from surgical procedures, the habituation to the OR arena occurred 7 days post-surgery, and the sample/reactivation/test phase 11 days after surgery.

### 2.3. Preparation and injections of drugs

DL-Propranolol was obtained from Sigma-Aldrich (MO/USA), prepared in 0.9% saline (NaCl), and injected into the hippocampus through the guide cannula employing a gingival needle connected to a Hamilton syringe (NV/USA). The drug infusion was performed at a speed of 0,5 µL/min and the needle was kept in place for 30-60 s to avoid drug reflux.

### 2.4. post-experimental confirmation of the injection sites

After behavioral experiments, animals were sacrificed (CO_2_ chamber, 10-30%) and immediately injected with methylene blue through the guide cannula (1 µL methylene blue). The brains were removed and the cannula placement was verified during sectioning using a cryostat (n = 27; Figure 1b). Infusions comprised most of the dorsal hippocampus, encompassing CA1, CA3, and dentate gyrus areas. All sampled animals had correct cannula placement.

### 2.5. Statistical analysis

The Shapiro-Wilk test was used to test the continuous variables against the normal distribution. Two-way ANOVAs were used to test the main effects of the object and treatment (propranolol infusion), followed by posthoc analysis using Tukey’s multiple comparisons tests. Data are expressed as Mean ± SEM, the main posthoc results are also expressed using confidence intervals (CI) and mean difference. The statistical analysis was conducted in GraphPad Prism v.9 and the statistical significance threshold was set at α = 0.05. The main effects are described in Table 1.

**Table 1.** Two-way ANOVA with Tukey’s multiple comparisons test.

| Propranolol dose |  | % of total variation | F (DFn, DFd) | P value |
| --- | --- | --- | --- | --- |
| Sample (5 $\mu\text{g}$) | Interaction | 0.5465 | $F(3, 246) = 0.8712$ | $P = 0.4566$ |
| | Object | 47.43 | $F(3, 246) = 75.62$ | **\*$P < 0.0001$** |
| | Treatment | 0 | $F(1, 246) = 0.000$ | $P > 0.9999$ |
| Reactivation (5 $\mu\text{g}$) | Interaction | 4.913 | $F(5, 112) = 1.782$ | $P = 0.1222$ |
| | Object | 33.29 | $F(5, 112) = 12.08$ | **\*$P < 0.0001$** |
| | Treatment | 0.000 | $F(1, 112) = 7.613\text{e-}031$ | $P > 0.9999$ |
| Sample (10 $\mu\text{g}$) | Interaction | 1.95 | $F(3, 264) = 3.574$ | $*P = 0.0146$ |
| | Object | 50.05 | $F(3, 264) = 91.76$ | **\*$P < 0.0001$** |
| | Treatment | 0 | $F(1, 264) = 0.000$ | $P > 0.9999$ |
| Reactivation (10 $\mu\text{g}$) | Interaction | 10.4 | $F(5, 118) = 6.073$ | **\*$P < 0.0001$** |
| | Object | 49.56 | $F(5, 118) = 28.95$ | **\*$P < 0.0001$** |
| | Treatment | 0.000 | $F(1, 118) = 6.130\text{e-}031$ | $P > 0.9999$ |

*\* $p < 0.05$ , in bold are $p < 0.001$*

## 3. Results

Previous studies have reported that systemic propranolol administration disrupts OR memory reconsolidation, evident in the increased exploration of a familiar object during the test phase (Liu et al., 2015; Villain et al., 2016). It is known that this systemic effect does not involve the basolateral amygdala, since it has been reported that intra-basolateral amygdala infusions do not disrupt OR memory reconsolidation (Maroun and Akirav, 2008). Here we address the involvement of the hippocampus by exploring whether intra-hippocampal infusions of propranolol disrupt OR memory reconsolidation.

During the sample phase, the post-hoc analysis indicates that animals of all groups equally explore both objects, A and B [Propranolol 5 µg experiment (treatment and saline): Mean difference -0.676, -5.212 — 3.859 (CI), p = 0.980; Propranolol 10 µg experiment (treatment and saline): Mean difference 3.815, -0.158 — 7.788 (CI), p = 0.065]. During the reactivation phase, animals of all groups explored significantly more the novel object C than the familiar object A, indicating successful OR memory consolidation [Figure 2a; Propranolol 5 µg experiment (treatment and saline): Mean difference -23.63, -31.20 — 22.05 (CI), p < 0.0001; Figure 3a; Propranolol 10 µg experiment (treatment and saline): Mean difference -25.21, -29.18 — 21.23 (CI), p < 0.0001].

**Figure 2.**
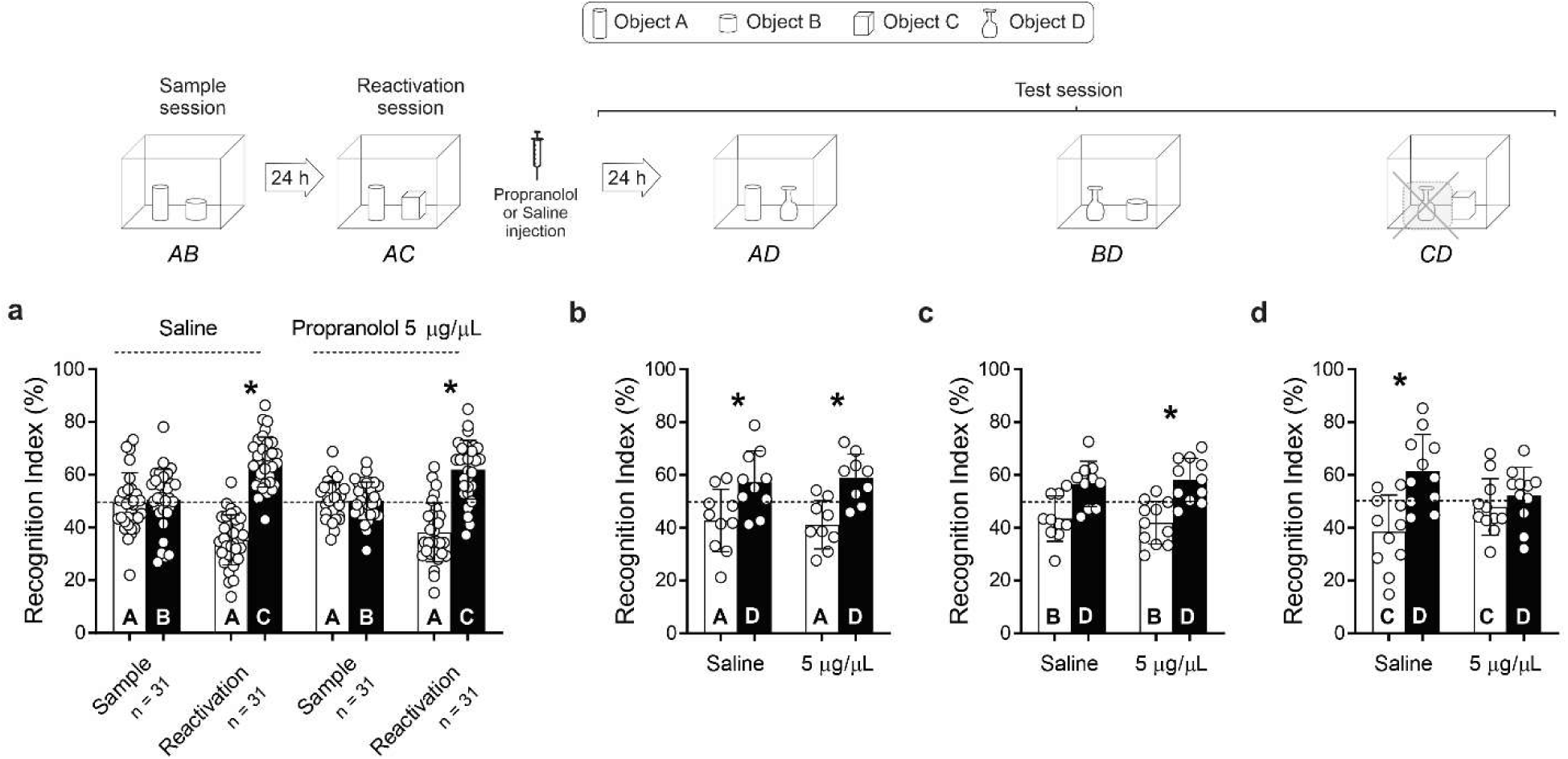
Blockage of the hippocampal β-adrenergic system by propranolol (5 μg/side) hampered the reconsolidation of a novel recognition memory trace. (**a**) Sample and reactivation phases: both experimental groups explored sample phase objects equally and were able to discriminate the novel object C at the reactivation phase. (**b-d**) The blockage of the hippocampal β-adrenergic system impaired the retention of the memory for the novel object C (Right panel; Saline, n = 11; Propranolol 5 μg/side; n = 11). Data are presented as the Mean ± SEM. *indicate significant difference, p < 0.05 (Tukey posthoc tests) – main effects are described in Table 1.

**Figure 3.**
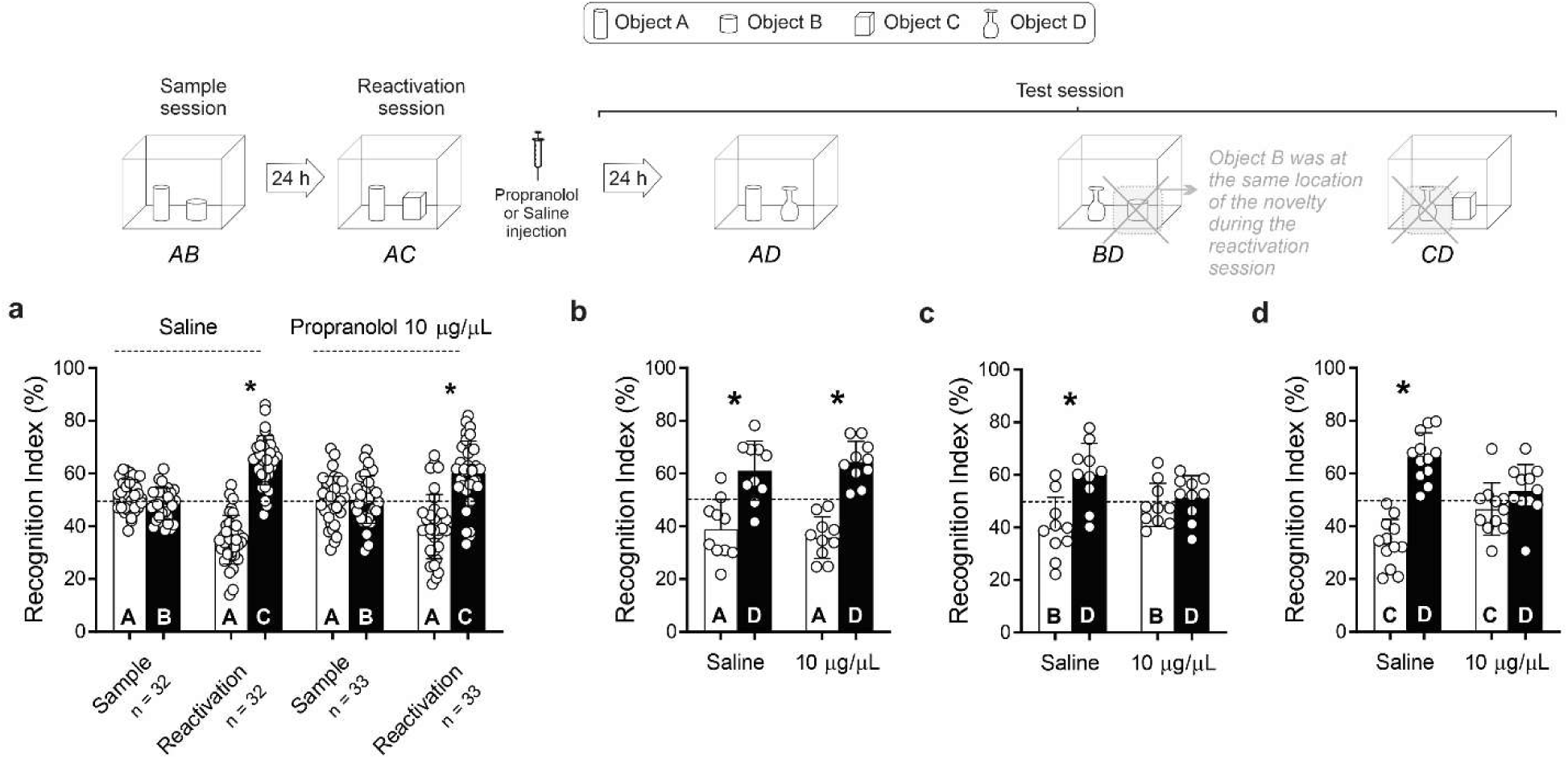
A high dose of propranolol (10 μg/side) affected the spatial component of a consolidated memory trace related to novel conjunctive events. **(a)** Similar to the previous experiment, in the sample phase both experimental groups explored sample phase objects equally and were able to discriminate the novel object C at the reactivation phase. **(b-d)** We confirmed the findings of the previous experiment: the blockage of the hippocampal β-adrenergic system impaired the retention of the memory for the novel object C (Right panel; Saline, n = 12; Propranolol 10 μg/side; n = 11). Additionally, infusion of propranolol (10 μg/side) hindered the memory of the familiar object (B) placed at the same position as the novelty during the reactivation phase (Middle panel; Saline, n = 10; Propranolol 10 μg/side; n = 11). Data are presented as the Mean ± SEM. *indicate significant difference, p < 0.05 (Tukey posthoc tests) – main effects are described in Table 1.

After reactivation, animals received intra-hippocampal infusions of saline or propranolol (5 µg or 10 µg/side) and the next day, were tested. During the test session, animals were exposed to a novel object, D, in combination with one of the following: the reactivated familiar object A; the non-reactivated familiar object B; object C, which was novel at the time of the reactivation. This allowed us to verify which memory component was affected by the treatments.

We first investigated the effect of a low and high dose of propranolol (5 µg/side and 10 µg/side) upon OR reconsolidation. When tested in the presence of the familiar object A (previously presented in both sample and reactivation sessions) and a novel object D; both saline-treated and propranolol-treated animals displayed a preference for the novel object [Figure 2b; Propranolol 5 µg experiment: Mean difference (treatment) -17.75, -32.24 — 3.26 (CI), p < 0.007 and Mean difference (saline) -14.58, -28.32 — 0.839 (CI), p < 0.031; Figure 3b; Propranolol 10 µg experiment: Mean difference (treatment) -31.06, -43.59 — 18.54 (CI), p < 0.0001 and Mean difference (saline) -22.16, -34.69 — 9.63 (CI), p < 0.0001]. This indicates that memory elements lacking novelty are not affected by intra-hippocampal propranolol administration.

When tested in the presence of object B - which was previously presented only in the sample phase (in the same spatial location of the novelty during the reconsolidation phase), and the novel object D. A similar outcome was observed, indicating that memory elements lacking novelty are not affected by a low dose of intra-hippocampal propranolol administration [Figure 2c; Propranolol 5 µg experiment: Mean difference (treatment) 16.24, 3.14 — 29.34 (CI), p = 0.006 and Mean difference (saline) -13.19, -0.55 — 26.93 (CI), p = 0.067]. These results of this first experiment also support the idea of a weaker memory for object B since saline-treated animals were slightly above the significance levels in this first experiment (p = 0.067); with no evident effects of 5 µg of propranolol infusion (p = 0.006). To confirm these findings, we performed an additional experiment using 10 µg of propranolol. In this experiment, the saline-treated animals were able to recognize object B (p < 0.0001), not evident in the group of animals treated with the high dose of propranolol (p = 0.961) [Figure 3c; Propranolol 10 µg experiment: Mean difference (treatment) 3.42, -8.51 — 15.37 (CI), p = 0.961 and Mean difference (saline) 20.53, 8.00 — 33.06 (CI), p < 0.0001]. Therefore, animals treated with a high dose of propranolol showed no preference for the novel object. This suggests that novelty can reactivate and destabilize memory elements that are weaker or indirectly associated by a spatial location. In order to disrupt an indirectly reactivated element, however, a higher dose of propranolol is required. This effect of high dose propranolol on memory components was evident in the interaction between treatment and object type - only evident with the high dose of propranolol (Propranolol 10 µg: F_objectXtreatment(5, 118)_ = 6.073, p < 0.0001 - Table 1).

When animals were tested with objects C (novel at the time of the reactivation phase) and D, there was a significant difference between groups [Figure 2d; Propranolol 5 µg experiment: Mean difference (treatment) 4.30, -8.79 — 17.41 (CI), p = 0.932 and Mean difference (saline) - 22.91, 9.81 — 36.02 (CI), p < 0.0001; Figure 3d; Propranolol 10 µg experiment: Mean difference (treatment) 7.80, -3.63 — 19.24 (CI), p = 0.362 and Mean difference (saline) 32.80, 21.37 — 44.23 (CI), p < 0.0001]. Post-hoc analysis indicated that while controls displayed a preference for the novel object D [saline: p < 0.0001 (5 µg experiment) and p < 0.0001 (10 µg experiment), propranolol-treated animals explored both objects equally [treatment: p = 0.932 (5 µg experiment) and p = 0.362 (5 µg experiment)]. This indicates that β-AR antagonism in the hippocampus blocks OR reconsolidation, thus preventing memory from updating with novel information (*i.e*., incorporating information of object C). The information that was already familiar at the time of reactivation (memory of object A), however, was not affected.

## 4. Discussion

Our results corroborate previous reports indicating that the intra-peritoneal administration of propranolol after a reactivation session (10 mg/kg) impairs object recognition reconsolidation in mice (Liu et al., 2015; Villain et al., 2016). It is known that propranolol crosses the blood-brain barrier and acts on multiple targets, such as the basolateral amygdala, the perirhinal cortex, and the hippocampus (McAinsh and Cruickshank, 1990). It has also been shown that intra-basolateral amygdala infusion of propranolol does not impair the reconsolidation of an object recognition memory (1.5µg/µL) (Maroun and Akirav, 2008). Our findings place the dorsal hippocampus as a major contributor to the systemic effect of propranolol on the reconsolidation of recognition memories.

The hippocampus mediates the addition of novel information to a previously consolidated memory trace (Haubrich et al., 2015; Rossato et al., 2007), a process that requires changes in synaptic strength. The β-ARs are key modulators of hippocampal synaptic plasticity (Hagena, Hansen & Manahan-Vaughan, 2016). As such, it has been demonstrated that hippocampal β-ARs inactivation disrupts memory reconsolidation and updating in several behavioral tasks (Dębiec et al., 2011; Gazarini et al., 2013; Muravieva and Alberini, 2010; Otis et al., 2013; Sara et al., 1999; Tronel, 2004). Our results go in line with these previous findings showing that hippocampal β-ARs are critical for the updating of OR memories. We also show that only the memory trace related to the novel object was destabilized since propranolol did not impair memory of a familiar object presented both during the sample and reactivation phases - regardless of dose. Nonetheless, recognition memory of an object not presented during the reactivation phase, but placed at the same position as the novelty (Object B), was hampered by higher doses of intra-hippocampal propranolol infusion. This suggests that elements of an OR memory can be indirectly reactivated and disrupted by propranolol. The novel OR protocol utilized in the present study has a strong spatial component (Rossato et al., 2007). Because distinct objects are presented during the sample phase, distinct spatial locations are acquired for objects A and B. Our results suggest that reconsolidation of the memory of an object can be achieved by presenting novel information at the same position where the familiar object was once located. In this way, novelty would trigger a destabilization process for the spatial component of this memory engram, therefore affecting elements linked to it. Indeed, it has been proposed that the hippocampal formation mediates such kind of conjunctive representations (Rudy et al., 2004) (Figure 4). The fact that only high doses of intra-hippocampal propranolol immediately after the reactivation session disrupt the reconsolidation of this memory trace will be discussed further below.

**Figure 4.**
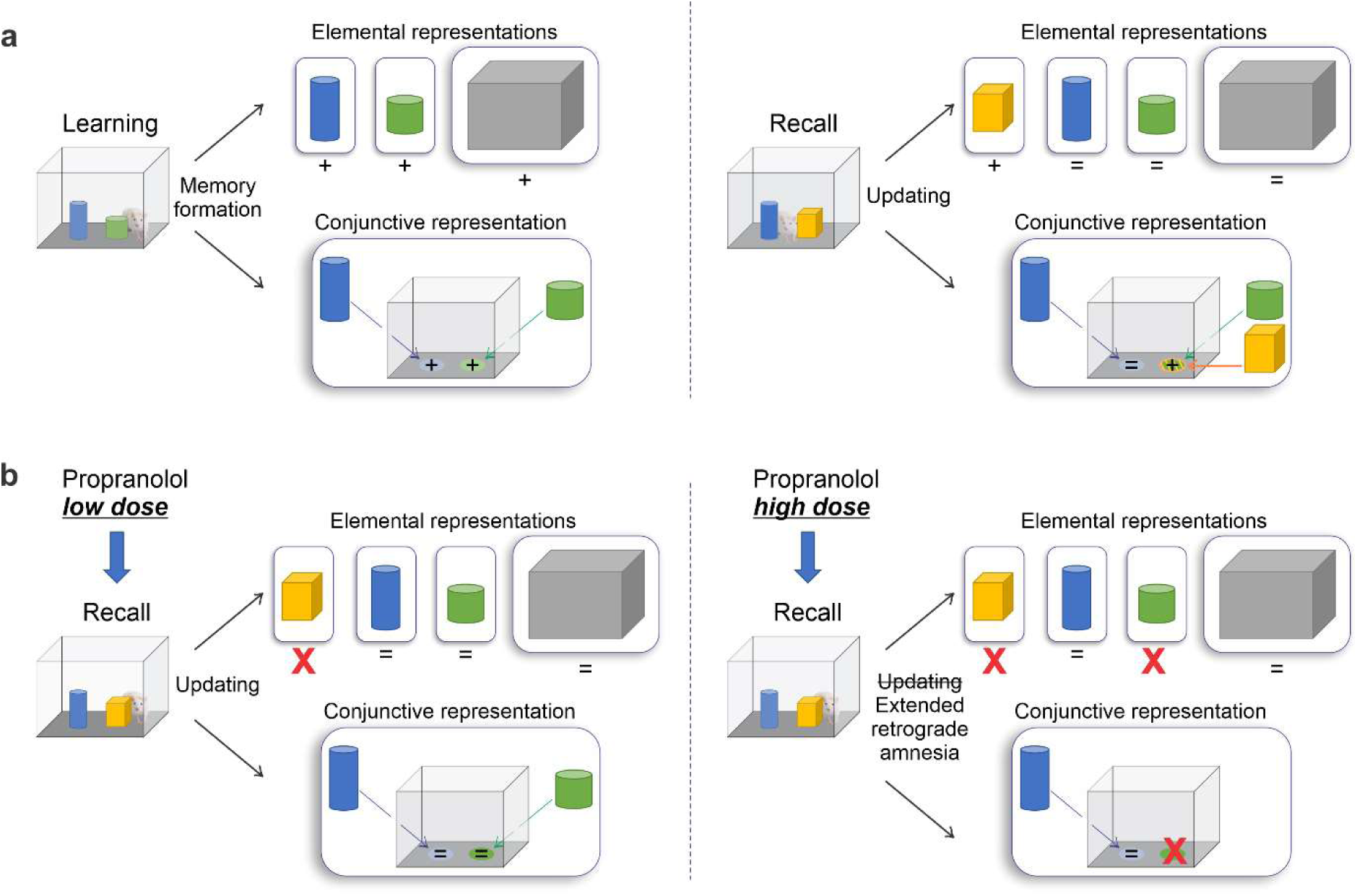
Two-process model: elemental and conjunctive components may be affected by propranolol, depending on the dose. (A) Schematics depicting the elemental (objects and arena) and conjunctive (spatial association between the objects and arena). At first, all components are new and compose the memory formation phase. A novel element is added (novel object) in a specific location and the learned memory trace is updated. (B) Intra-hippocampal infusion of propranolol immediately after recalling and updating the memory trace hampers the reconsolidation of the initial memory trace during recall. Nonetheless, recognition memory of an object not presented during the reactivation phase, but placed at the same position as the novelty, was also hampered by higher doses of intra-hippocampal propranolol infusion. In the latter, both elemental and conjunctive representations are affected. The model was based on (Rudy et al., 2004).

The hippocampal formation is typically associated with the spatial component of recognition memories, with specific cell types encoding place-object associations (Vann and Albasser, 2011). On the other hand, perirhinal cortices would account mostly for the sensorial information of the object, such as geometric shapes, lights, visual-tactile stimulus (Kealy and Commins, 2011). The perirhinal cortices are the major sensorial input to the hippocampus (Brown and Xiang, 1998). Notwithstanding the specific roles, the interaction between the perirhinal cortex and the hippocampus is required for recognition memory (Warburton and Brown, 2010). Nonetheless, the perirhinal cortex does not provide significant excitatory inputs to the dentate gyrus or CA1 when compared to other afferents, *e.g*., the lateral perforant path, lateral entorhinal cortex, and amygdala-entorhinal transition (Canning and Leung, 1998). Indeed, the entorhinal cortex projects strongly to all regions of the hippocampal formation, whereas the perirhinal and postrhinal cortices project weakly and only to CA1 and subiculum (Agster and Burwell, 2013). In other words, the entorhinal cortex has a major role in spatial codification via strong, intricate, reciprocal connectivity with the hippocampus (Morrissey and Takehara-Nishiuchi, 2014). Therefore, the sensorial afferents arriving from the perirhinal cortex are more localized and less redundant and, thereby, more susceptible to the effects of intra-hippocampal injections.

Interestingly, lesions to the medial entorhinal cortex preferentially impair the recognition of the spatial arrangement of objects relevant to the spatial location of an experience (Morrissey and Takehara-Nishiuchi, 2014). The rat hippocampus is vital for the accurate performance of tests of allocentric spatial memory (Aggleton et al., 2000) and multiple factors could contribute to our findings. Given the pivotal importance of the perirhinal cortex on the ‘familiarity discrimination’ component of recognition memories, further studies should investigate the effects of intra-cortex perirhinal propranolol infusion on recognition memory reconsolidation.

## 5. Conclusion

To our knowledge, the present study is the first to provide information related to the effects of intra-hippocampal propranolol infusions on OR memory reconsolidation. The addition of novel information to a previously consolidated reactivated memory was impaired by propranolol administration. In higher doses, propranolol affected the spatial component of the memory trace, evident in the difficulty in recognizing a familiar object placed where the novelty was presented during the reactivation session. This indicates that both elemental and conjunctive components of OR memory may be affected by propranolol, depending on its dose, and this effect was specific since memory elements not linked to the addition of novel information were spared. The findings of this study corroborate recent work on reconsolidation of recognition memories and systemic administration of propranolol in mice (Villain et al., 2016). Our results place the hippocampal formation as a major contributor to the effects of propranolol administration upon OR memory reconsolidation. Taking together, the present outcomes shed light on the effects of propranolol on non-targeted memories during PTSD treatment by revealing that both elemental and conjunctive associations can be disrupted by post-reactivation propranolol depending on its dose. Further clinical studies should examine possible recognition memory deficits during propranolol PTSD treatment.

## Acknowledgments

This study was supported by Coordenação de Aperfeiçoamento de Pessoal de Nível Superior.

## Data Availability

The data that support the findings of this study are available from the corresponding author upon reasonable request.

